# Chromatin-informed inference of transcriptional programs in gynecologic and basal breast cancers

**DOI:** 10.1101/333757

**Authors:** Hatice U. Osmanbeyoglu, Fumiko Shimizu, Angela Rynne-Vidal, Petar Jelinic, Samuel C. Mok, Gabriela Chiosis, Douglas A. Levine, Christina S. Leslie

## Abstract

Epigenomic data on transcription factor occupancy and chromatin accessibility can elucidate the developmental origin of cancer cells and reveal the enhancer landscape of key oncogenic transcriptional regulators. However, in many cancers, epigenomic analyses have been limited, and computational methods to infer regulatory networks in tumors typically use expression data alone, or rely on transcription factor (TF) motifs in annotated promoter regions. Here, we develop a novel machine learning strategy called PSIONIC (patient-specific inference of networks informed by chromatin) to combine cell line chromatin accessibility data with large tumor expression data sets and model the effect of enhancers on transcriptional programs in multiple cancers. We generated a new ATAC-seq data set profiling chromatin accessibility in gynecologic and basal breast cancer cell lines and applied PSIONIC to 723 RNA-seq experiments from ovarian, uterine, and basal breast tumors as well as 96 cell line RNA-seq profiles. Our computational framework enables us to share information across tumors to learn patient-specific inferred TF activities, revealing regulatory differences between and within tumor types. Many of the identified TF regulators were significantly associated with survival outcome in basal breast, uterine serous and endometrioid carcinomas. Moreover, PSIONIC-predicted activity for MTF1 in cell line models correlated with sensitivity to MTF1 inhibition. Therefore computationally dissecting the role of TFs in gynecologic cancers may ultimately advance personalized therapy.

## INTRODUCTION

Cancers arise through the accumulation of genetic and epigenetic alterations that lead to widespread gene expression changes. Transcription factors (TFs) are instrumental in driving these gene expression programs, and the aberrant activity of TFs – induced downstream of activated oncogenic signaling or in concert with epigenetic modifiers – often underlies the altered developmental state of cancer cells and acquisition of cancer-related cellular phenotypes. Data-driven computational strategies may help to infer patient-specific transcriptional regulatory programs and to identify and therapeutically target the TFs that lead to cancer phenotypes. Ultimately, such strategies could be used to personalize therapy and improve patient outcomes.

While several successful methods have been proposed for learning patient-specific regulatory programs, most regulatory network inference approaches in cancer use expression data only [1] or at best rely on analysis of transcription factor (TF) motifs in annotated promoter regions [2-4]. However, in a few cancers — notably luminal breast and prostate cancer — ChIP-seq analyses of key transcriptional regulators, estrogen receptor (ER) and androgen receptor (AR) respectively, in both cell line models and tumors have revealed the importance of enhancers distal to gene promoters in gene regulatory programs. Incorporating DNA sequence information at intronic and intergenic enhancers should therefore improve the modeling of transcriptional regulation in tumors. Leveraging epigenomic data from cell line models, while imperfect, provides a feasible means to make a potentially large advance in the computational dissection of dysregulated gene expression programs in tumors.

Extensive pan-cancer genomic analyses have shown that the same genes and pathways are targeted by somatic alterations across multiple tumor types. These results suggest that pan-cancer modeling of regulatory programs could also be informative, as similar TFs may be dysregulated across cancers. So far, however, methods for inferring patient-specific regulatory programs have been applied to one cancer type at a time [1, 5]. *Multitask learning* (MTL) refers to machine learning algorithms that learn models for different problems that share information and/or parameters and provides a statistical framework for learning patient-specific regulatory models across multiple cancers [6]. MTL can improve accuracy by making use of limited data (small sample sizes) in each task by sharing information through the common model. This is especially important when reconstructing regulatory networks from high-throughput data because the number of parameters to fit is very large relative to the number of samples. In addition, extensive training data from more common tumor types may be able to compensate for smaller sample sizes in similar but rarer cancers.

Large-scale cancer genomics projects such as The Cancer Genome Atlas (TCGA) and others have suggested molecular similarities between gynecologic cancers from different sites of pelvic origin and breast cancers. Specifically, uterine serous carcinomas (UCS), high-grade serous ovarian carcinomas (HGSOCs), and triple negative breast cancers (TNBCs) share frequent *TP53* somatic mutations and widespread somatic copy number alterations. HGSOCs and TNBCs also both display inactivation of similar DNA repair pathways. Though each gynecologic disease has a variety of histologic subtypes, the most common and aggressive tumors including high-grade serous ovarian carcinomas (OV), UCS, and the serous-like subset of uterine (UCEC) as well as basal breast cancer were studied by TCGA. These tumors all lack adequate treatment options for recurrent disease and accurate predictors of response and resistance. Inferring patient-specific transcriptional regulatory programs may identify and eventually enable therapeutic targeting of transcriptional mechanisms underlying gynecologic malignancies for individualized treatment.

To improve inferring regulatory programs across cancer types, we developed PSIONIC (patient-specific inference of networks incorporating chromatin), a multitask learning method that jointly models transcriptional networks for several related cancer types by leveraging chromatin accessibility data in representative cancer cell lines. More specifically, PSIONIC integrates regulatory sequence from ATAC-mapped promoters and enhancers from a panel of cancer cell lines with RNA-seq data from patient tumors in order to infer patient-specific TF regulatory activities. We applied our approach to 723 RNA-seq experiments from gynecologic and basal breast cancer tumors as well as 96 cell lines, using a novel ATAC-seq data set for cell line models of these cancers. ATAC-seq data from cell lines allowed us to incorporate DNA sequence information at intronic and intergenic enhancers to improve the modeling of transcriptional regulation from tumor data. Although much work has been done in regression-based inference of transcriptional regulation from *cis* regulatory information in a single tumor type, we use multitask learning across different tumor types to jointly learn patient-specific regulatory model. Our analysis identified key transcriptional regulators as well as new prognostic markers and therapeutic targets.

## RESULTS

To systematically identify TFs that drive tumor-specific gene expression patterns across multiple cancer types, we developed the PSIONIC computational framework (**Figure 1**). We start with an atlas of chromatin accessible events derived from cell line models of the tumor types to be analyzed, using ATAC-seq profiling data (**Methods**). We represent every gene by its feature vector of TF binding scores, where motif information is summarized across all promoter, intronic, and intergenic chromatin accessible sites assigned to the gene (**Methods**). Single-task learning of a patient-specific regulatory model simply learns the TF activities that predict normalized gene expression levels in each tumor independently, using regularized regression (**Figure 1A**, **Methods**). In PSIONIC, we instead adopt a multitask learning approach called GO-MTL [7] to represent patient-specific TF activity model vectors across multiple tumor types as linear combinations of *latent regulatory programs*, where both the coefficients in the linear combination and the latent models are learned jointly by regression against all the normalized tumor expression profiles (**Figure 1B**, **Methods**). The latent regulatory programs capture common TF-gene regulatory relationships across patients both within and between tumor types.

**Figure 1.**
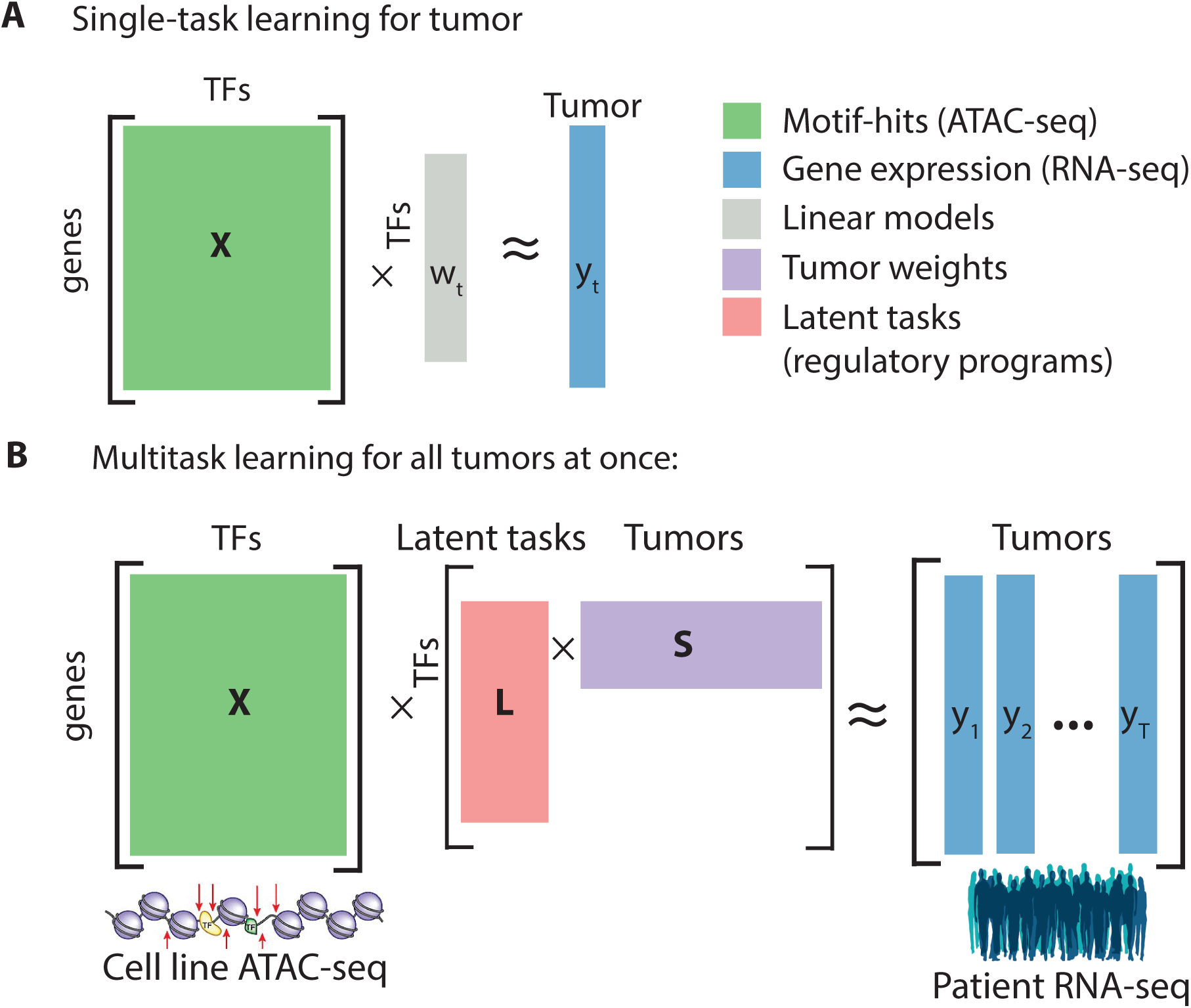
Overview of PSIONIC algorithm. The input to our framework includes ATAC-seq profiles, TF recognition motifs, and tumor expression datasets. A schematic comparison of **(A)** single task learning and **(B)** multitask learning models.

### ATAC-seq analyses of gynecologic and basal breast cancer cells

To enable PSIONIC modeling for gynecologic and basal breast tumors, we first generated a reference chromatin accessibility atlas for uterine (endometrioid, serous, carcinosarcoma), ovarian serous, and basal breast cancers using a panel of twelve cancer cell lines representing these five tumor types using the assay for transposase-accessible chromatin with high-throughput sequencing (ATAC-seq). We first performed peak calling on ATAC-seq profiles, using two biological replicates per cell type to control for irreproducibility discovery rate (IDR), and assembled an atlas of ∼282K reproducible accessibility regions for all cell lines as well as tumor type specific atlases ranging from ∼93K to ∼153K reproducible regions **(Supplementary Table 1, Methods)**. Principal component analysis (PCA) identified heterogeneity in the chromatin accessibility landscape in these gynecologic and basal breast cancer cell lines (**Figure 2A)**. Notably, ovarian and basal breast cancer cell lines displayed more similar chromatin accessibility profiles compared to most of the uterine cancer cell lines. Interestingly, for the two uterine carcinosarcoma cell lines, the copy number high SNU685 cell line clustered with ovarian and basal breast cancer cell lines, whereas JHUCS1 clustered with uterine endometrioid cell lines.

**Figure 2.**
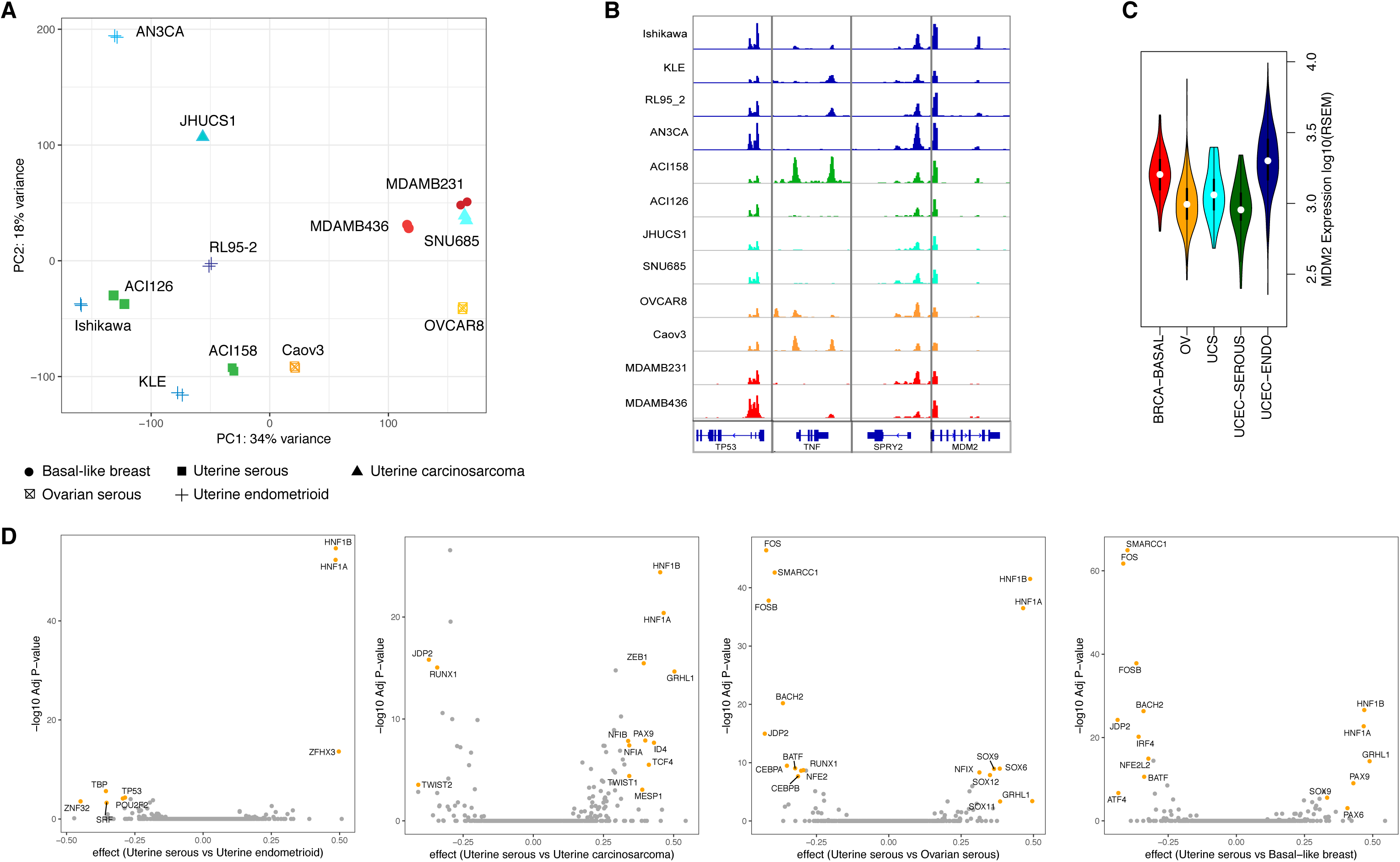
ATAC-seq analysis identifies key TFs in gynecologic and basal breast cancer cells. **(A)** Unsupervised principal component analysis based on the chromatin accessibility for all 12 cell lines at each of the 10K most variable chromatin accessible regions in the cell line panel. Samples are color coded according to their tumor type of origin. Each symbol represents a single biological replicate. **(B)** Normalized ATAC-seq profiles at important genes. Profiles represent the union of all biological replicates for each cell type. Genomic coordinates for the loci: *TP53*, chr17:7571720-7590868; *TNF*, chr6:31543344-31546112; *SPRY2*, chr13:80910112-80915086; *MDM2*, chr12:69201952-69239324. All *y*-axis scales range from 0-10 in normalized arbitrary units. The *x*-axis scale is indicated by the scale bars. **(C)** Violin plots indicate the distribution of *MDM2* gene expression across tumor types. MDM2 gene expression is higher in uterine endometrioid carcinomas (UCEC-ENDO) compared to other tumor types. **(D)** Pairwise comparison of transcription factor motifs enriched in differentially accessible regions in cell lines. Volcano plot showing effect size versus –log10(*P*), using a Bonferroni correction to adjust *P* values for each plot. TF symbol annotations are written where the absolute value of the effect size is in **(A)** at least top 30 and adjusted *P* < 10^−3^. The foreground occurrence is the number of peaks containing a particular TF motif within the group of 5000 upregulated or 5000 downregulated peaks according to log2 fold-change read counts, respectively. The background occurrence is the number of peaks containing a particular TF motif found among all the differentially accessible peaks.

Next, we assigned each accessible region in the tumor type specific atlas to the nearest gene **(Figure 2B)** and we defined the regulatory locus complexity of a gene [8] as the total number of accessible regions within the tumor type. More specifically, we grouped genes into three equally sized classes (tertiles) on the basis of their regulatory complexity in tumor type specific atlases. Complexity classes were defined by dividing genes at the 33rd and 66th percentiles of the distribution of the number of accessible regions to produce groups with similar numbers of genes. We found that the normalized expression levels of low-complexity genes were lower than high-complexity and medium-complexity genes in tumor samples from TCGA for each tumor type (*P* < 1 x 10^−16^, one-sided Kolmogorov-Smirnov (KS) test for all comparisons). This observation is illustrated by the region surrounding the *MDM2* gene. Despite the ubiquitous accessibility of the MDM2 promoter, nearby distal regulatory elements of MDM2 were more accessible in uterine endometrioid cell lines, consistent with higher *MDM2* gene expression observed in corresponding tumor samples from the TCGA cohort (**Figure 2C**).

### Motifs underlying differential accessibility in cell lines

We next sought to determine the TFs that are most associated with open chromatin for each tumor type through motif analyses and differential accessibility. First, we performed motif analysis in each chromatin accessible regions in the common atlas (**Methods**). Next, we examined the patterns of gain or loss of chromatin accessible regions between each pair of tumor types by performing pairwise differential read count analysis on accessible regions. The heatmap in **Supplementary Figure 1** shows the patterns of differential accessibility found among ∼40,000 peaks across cell lines. Many TFs whose motifs were identified at differentially accessible regions between pairs of tumor types have roles in tumorigenesis (**Figure 2D**, **Supplementary Figure 2**). For example, chromatin peaks with HNF1 family motifs were more accessible in the endometrioid subset of uterine cell lines compared to others (*P* < 1 x 10^−16^, one-sided Kolmogorov-Smirnov (KS) test). HNF1β is associated with cancer risk in several tumors, including hepatocellular carcinoma, pancreatic carcinoma, renal cancer, ovarian cancer, endometrial cancer, and prostate cancer [9]. KLF and ETS family motifs were more accessible in the endometrioid subset of uterine cell lines as well as ovarian serous cell lines in contrast to others (*P* < 1 x 10^−16^, one-sided Kolmogorov-Smirnov (KS) test). These TFs have been implicated in the pathogenesis of these endocrine-responsive cancers of female reproductive tissues [10, 11]. Chromatin peaks with FOS family motifs were more accessible in basal breast, ovarian serous and uterine carcinosarcoma and were less accessible in uterine endometrioid cell lines (*P* < 1 x 10^−16^, one-sided Kolmogorov-Smirnov (KS) test). FOS family TFs have been implicated as regulators of cell proliferation, differentiation, and transformation. While many identified TFs are known to play a role in other cancers, their impact on gene regulation has not been characterized in gynecologic and basal breast cancers. We therefore developed a regression framework to model the regulatory role of TFs on gene expression in tumor samples.

### Multitask regression explains gene expression profiles in tumor samples

We next used a multitask learning strategy across tumor types to learn patient-specific regression models to predict tumor gene expression from gene regulatory sequence derived from cell line ATAC-seq data. Our method assumes that observed gene expression levels in each tumor can largely be explained by the unobserved activities of a smaller number of TF regulatory proteins through correlation with TF binding motif scores. Moreover, our approach shares information across tumor samples and tumor types by representing each patient-specific regulatory model as a linear combination of a *latent regulatory models.*

Formally, we developed PSIONIC, a multitask-learning framework for integrating regulatory elements for each gene based on motifs in ATAC-mapped promoters and enhancers from cancer cell lines (**X**) with RNA-seq data from patient tumors (**Y**) in order to infer patient-specific TF regulatory activities (**W** = **LS**) (**Figure 1B**). More specifically, we adopted an algorithm for learning *grouping and overlap structure in multitask learning* (GO-MTL) [7]; here, the model does not assume a disjoint assignment of tasks (patients) to different groups (e.g. tumor type) but rather allows patient-specific models to overlap with each other by sharing one or more latent basis tasks, or *latent regulatory programs*. Here, the matrix **L** represents these latent regulatory programs, while **S**, the tumor weight matrix, captures the grouping structure and specifies the coefficients of the linear combination of latent regulatory programs for each tumor. Multitask learning enables selective sharing of information across other tumors, while standard single-task learning (STL) trains a regression model for each tumor independently.

We applied our approach to 723 uterine, ovarian, and basal breast tumors from TCGA in order to identify key TFs as potential common or cancer-specific drivers of expression changes. Our expression dataset included samples from five different tumor types, namely basal breast (BRCA-BASAL, n=92), high-grade serous ovarian (OV, n=255), uterine carcinosarcoma (UCS, n=57), uterine endometrioid carcinomas (UCEC-ENDO, n=272), and uterine serous carcinomas (UCEC-SEROUS, n=47). As motif data, we used binding site predictions for 352 human sequence-specific TFs based on motif hits from the CisBP database (**Methods**).

We first evaluated performance of PSIONIC as well as STL based on ridge regression for each tumor type (**Figure 3A**). For statistical evaluation, we computed the mean Spearman correlation between predicted and measured gene expression profiles on a validation set of held-out genes for each tumor type. We obtained significantly better regression performance than a STL approach based on ridge regression (*P* < 10^−8^, one-sided Wilcoxon signed-rank test). Similarly, our models with motif data from promoter and enhancer regions outperformed models where only motif hits in promoter regions were used (*P* < 10^−16^, one-sided Wilcoxon signed-rank test). In contrast, if we randomized motif hits for each chromatin accessible region across all motifs, or if we randomized accessible regions for each motif, then assigned to the nearest gene, the prediction performance also significantly decreased (*P* < 10^−16^, one-sided Wilcoxon signed-rank test).

**Figure 3.**
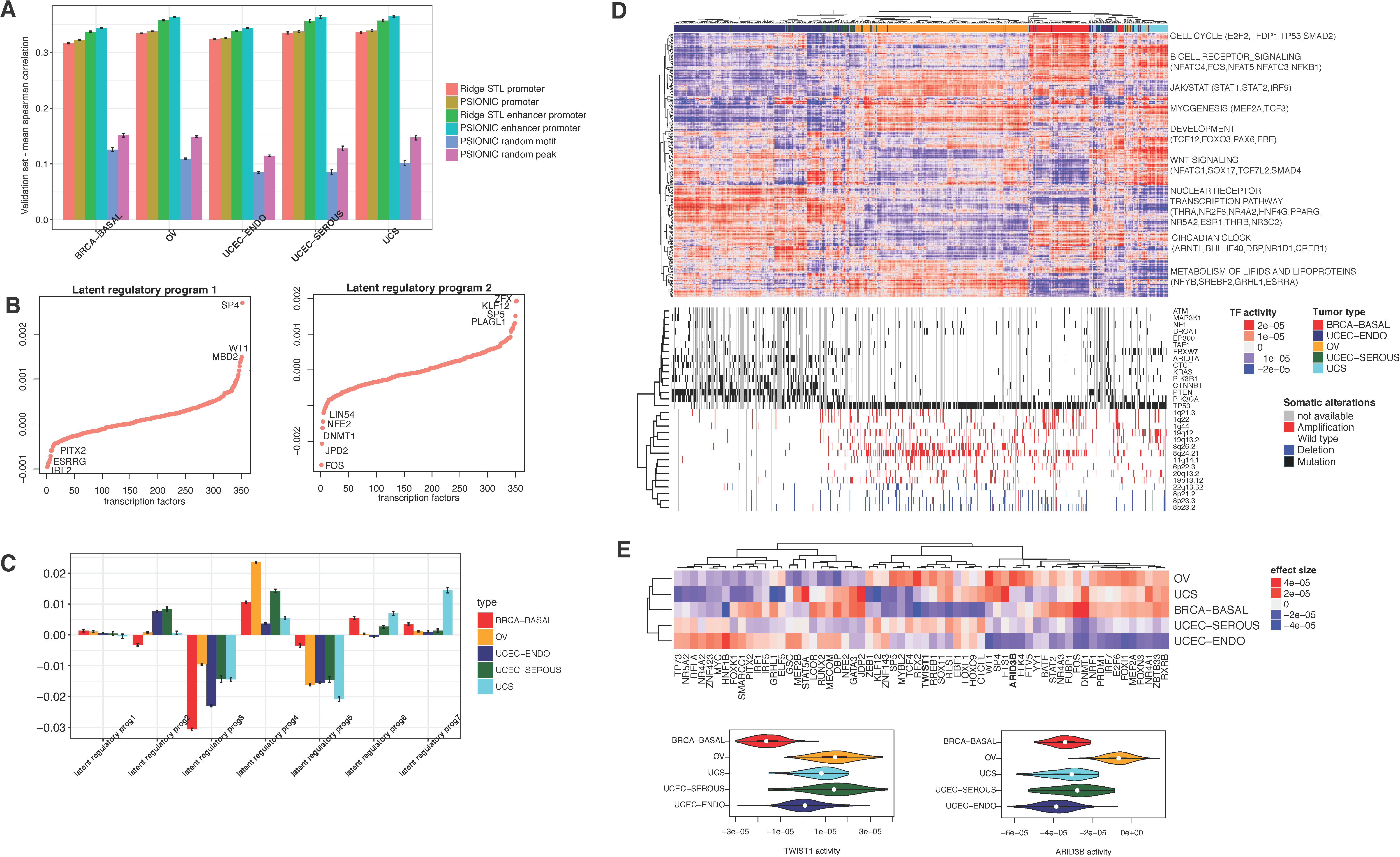
PSIONIC identifies regulatory features of tumor types. **(A)** Performance comparison of ridge regression and multitask learning models. Models built using TFs mapped to promoters as well as enhancers as features perform significantly better than those using TFs mapped to only promoters (*P* < 10^−16^, one-sided Wilcoxon signed-rank test); multitask learning models performed significantly better than single task learning models (*P* < 10^−8^, one-sided Wilcoxon signed-rank test). When the motif hits for each peak were randomized across TFs, or when peaks for each motif were randomized, and then peaks were assigned to the nearest gene, the performance of the models significantly decreased (*P* < 10^−16^, one-sided Wilcoxon signed-rank test). **(B)** Example of latent regulatory programs. TFs are ranked based on the magnitude of coefficients. **(C)** Average tumor weight matrix (**S**) grouped according to tumor type for each latent regulatory program. **(D)** Hierarchical clustering model vectors (**W**) on the TCGA tumor data sets. **(E)** Heatmap shows the mean inferred TF activity differences between samples in a given tumor type vs. those in all other tumor types. For each comparison, the absolute value of the mean inferred TF activity differences (effect sizes) are ranked and the union of top 20 TFs for each comparison are shown in the heatmap. Violin plots indicate the distribution of inferred ARID3B and TWIST1 TF activities across tumor types.

We found that the number of latent regulatory programs giving the best performance was k = 7 (**Figure 3B, Supplementary Figure 3**). **Figure 3C** shows a summary of mean tumor weights (**S**) across each tumor type for each latent regulatory program. For example, latent regulatory program 1 appears to capture a common gene regulatory program shared across all cancer types, whereas latent regulatory program 2 captures TF-gene regulatory relationships shared by uterine serous and endometrioid tumors. Hierarchical clustering of tumors by inferred TF activities, **W**=**LS**, as derived from the model largely recovered the distinction between the major tumor types (**Figure 3D**). In particular, serous endometrioid tumors have distinct patterns of TF activities, consistent with their differing expression and mutational patterns.

### Clinical outcome based on inferred TF activities

To investigate the clinical relevance of TF activities, we examined whether inferred TF activities were associated with therapeutic response. The standard of care for ovarian serous patients is aggressive surgery followed by platinum/taxane chemotherapy. After therapy, platinum resistant cancer recurs in approximately 25% of patients within six months [12]. Due to the clinical significance of this problem, we sought to determine TFs linked to platinum resistance in OV. Inferred TF activities of 7 TFs were significantly associated with platinum response including HIF-1α and ZNF423 (*t*-test, *P* < 0.05, Supplementary Figure 4). Consistent with our findings, HIF-1α has been associated with platinum resistance in a variety of cancers, including ovarian [13]. Moreover, ESR1 and ZNF423 have role in cancer cell proliferation [14, 15] and were significantly associated with platinum-sensitive tumors.

Next, we examined whether inferred activities of TFs were linked to survival data from the TCGA. The patient survival data and matched TF activities enabled us to perform TF-centric survival analyses to identify prognostic TFs within tumor type (TFs with FDR-adjusted *P* < 0.05, univariate Cox analysis). Many TFs had significant associations with survival outcome in BRCA-BASAL, UCEC-SEROUS and UCEC-ENDO (**Figure 4A** and **Supplementary Table 2**). For some TFs, the prognostic value has previously been reported in the literature; for example, PGR [16] has been associated to survival in uterine cancer. However, most of the identified prognostic TFs lack prior reports of a link to survival in these cancers, making them potential candidates for follow-up studies. For example, ETV6 inferred activity separated patients into high-and low-risk groups in UCEC-SEROUS (FDR < 0.05, univariate Cox analysis). ETV6 exhibits antitumor effects suppressing proliferation and metastatic progression in prostate cancer [17]. However, its role in uterine serous cancer has not been studied. A representative image of immunofluorescence staining on a primary uterine serous tumor shows protein level nuclear localization of ETV6 in tumor cells (**Figure 4B)**.

**Figure 4.**
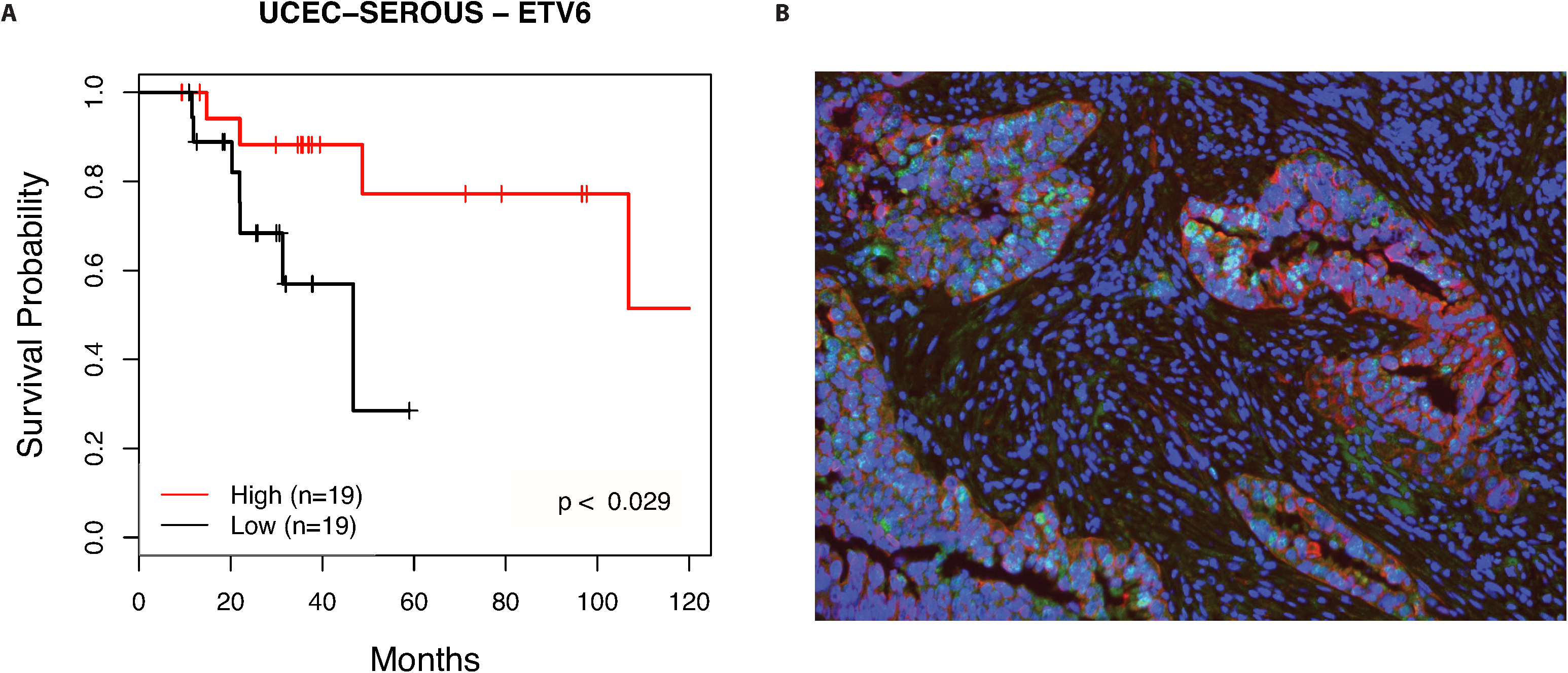
Identification of prognostic TFs based on inferred activities coupled with clinical survival. **(A)** Kaplan-Meier plot for UCEC serous patients stratified by the inferred activity of a ETV6, with patient numbers for each group in parentheses: patients with ETV6 activities are above mean, “High,” lower than mean, “Low”; *P <* 0.029, log-rank test. **(B)** Representative image of immunofluorescence staining on a primary endometrial serous tumor. Double staining for ETV6 (green) and cytokeratin (red), shows nuclear localization of ETV6 in tumor cells; DAPI (4’,6-diamidino-2-phenylindole): blue.

### Multitask regression identifies tumor type-specific TFs associated with expression

Next, we assessed TF-tumor type associations by t-test to compare inferred TF activity between samples in a given tumor type vs. those in all other tumor types. We corrected for FDR across TFs for each such pairwise comparison and identified significant TF regulators. Results are shown in **Supplemental Table 3**, **Figure 3E**. FUBP1, which regulates *c-Myc* gene transcription, had significantly higher inferred activity in BASAL-BRCA compared to gynecologic tumors; ARID3B activity was significantly higher in OV, consistent with its role in promoting ovarian tumor development, in part by regulating stem cell genes [18]; NR5A2 (also known as liver receptor homolog-1, LRH-1) was significantly higher in uterine endometriod tumors, consistent with its function in regulating metabolism and hormone synthesis. Also in agreement with previous reports, WT1 activity was significantly higher in ovarian serous [19] and uterine sarcoma [20]; TWIST1, a central player in the EMT, had increased activity in ovarian serous [21] and uterine serous cancers; YY1, which regulates various processes of development and differentiation and is involved in tumorigenesis of breast and ovarian cancer [22], had increased activity in these cancers.

Besides confirming key TFs from previous studies, our analysis also predicted novel TF regulators in gynecologic and basal breast cancers. For example, MEF2A, a transcriptional regulator implicated in muscle development, cell growth control, and apoptosis, had significantly higher activity in OV, BRCA-BASAL and UCS; the activity of MITF, microphthalmia-associated transcription factor, was significantly higher in uterine carcinosarcoma and displayed high variation across patients. Indeed, UCSs are characterized by an admixture of at least two histologically distinct components, one resembling carcinoma and another resembling sarcoma [23]. The roles of MEF2A and MITF have not been previously characterized in these cancers and may present promising targets for study and potentially for therapeutic intervention.

To investigate the potential of using PSIONIC-inferred TF activities to predict sensitivity to TF-targeted therapeutics, we decided to translate our model into cancer cell lines where drug sensitivity can be experimentally determined. Therefore, we first assembled a collection of basal-like, ovary and endometrium transcriptional profiles of immortalized human cancer cell lines from the CCLE [24], trained a PSIONIC model on this data set, and hence inferred cell line-specific TF regulatory activities. Regulatory models for cell lines to some extent recapitulated patient-specific tumor regulatory models (**Supplementary Figure 5**). While there are few drugs that directly target transcription factors, we were able to use the MITF inhibitor LOR-253 for a proof of principle analysis. We assessed our original panel of 10 cell lines for sensitivity to LOR-253 by measuring growth rate inhibition. Consistent with expectation, cell lines with higher inferred MTF1 activity had a greater decreased in growth rate after the treatment with LOR-253 (**Figure 5**). Overall, MTF1 inferred activity was significantly associated with growth rate inhibition by Spearman correlation analysis (*P* < 10^−2^ for these cell lines).

**Figure 5.**
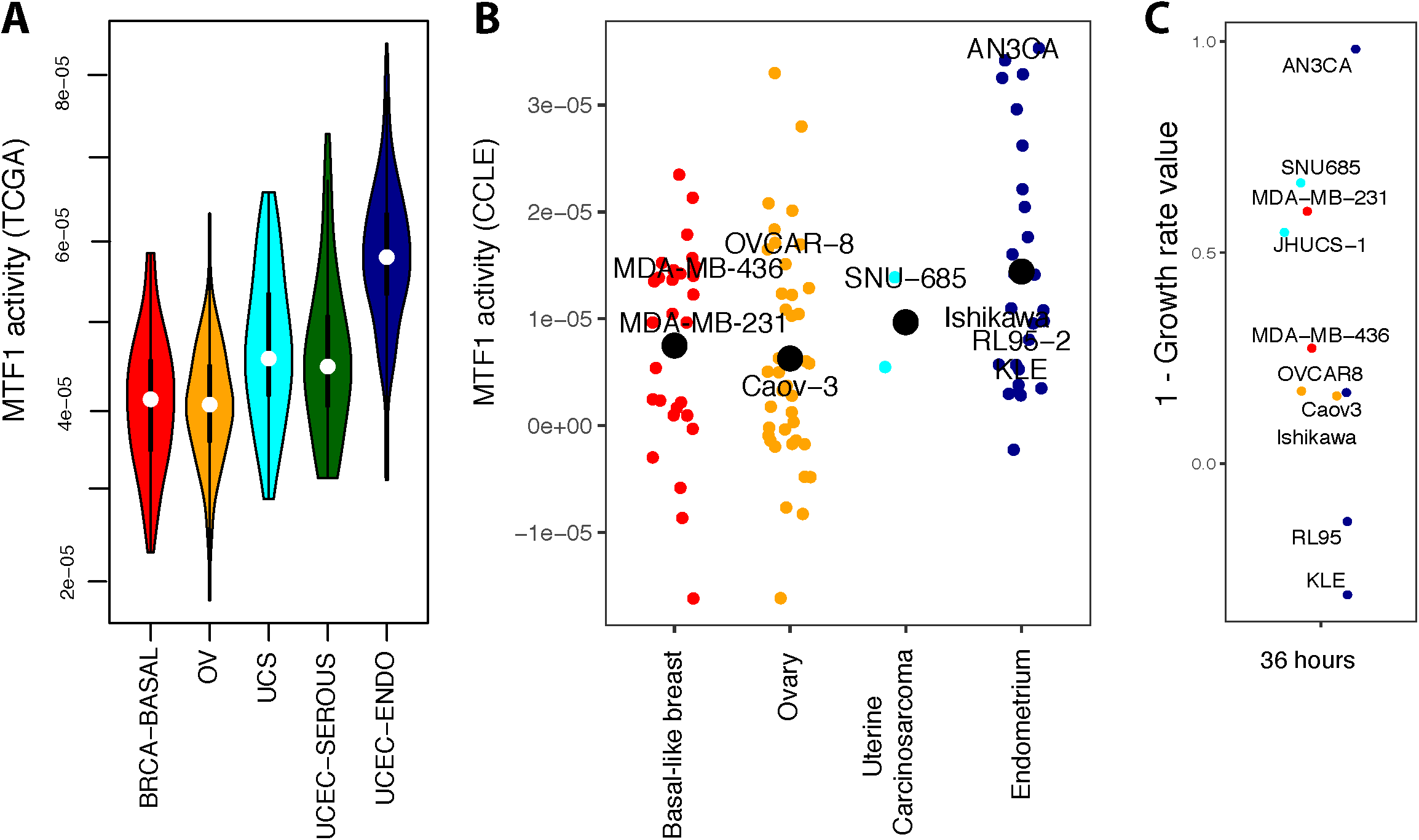
PSIONIC predicts cell line sensitivity to TF-targeted therapy. **(A)** Violin plots indicate the distribution of inferred MTF1 TF activities across tumor types. **(B)** We trained a PSIONIC model on 96 cell lines from the CCLE study. The dot plots show inferred MTF1 activities in basal-like breast, endometrium, ovary and uterine carcinosarcoma cell lines. **(C)** 1 – growth rate (GR) values [37] for MTF1 inhibitor LOR-253 36 hours after the treatment in gynecologic and basal breast cancer cell lines.

## DISCUSSION

With the development of high-throughput sequencing technologies, transcriptomic, proteomic, genomic profiles of tumor samples have been rapidly generated for diverse cancer types. Identifying differentially expressed genes or recurring mutations does not always clarify the molecular pathways that actually regulate tumor state and survival. There is still a large methodological gap between generating molecular profiles of tumor samples and understanding the molecular mechanisms underlying tumorigenesis and response to therapy.

Our PSIONIC method provides a systematic framework for integrating resources on regulatory genomics with tumor expression data to better understand gene regulation in cancers and infer patient-specific TF networks. PSIONIC uses a reduced rank representation model based on latent tasks, which helps regularize patient-specific regression models in light of noisy tumor gene expression data while sharing information between tumors and tumor types. Joint inference of TF activities across different tumor types may also reveal clinically relevant patient subgroups common to multiple cancers. As new ATAC-seq technologies for frozen tissue are developed [25], ATAC-seq will become feasible in clinical samples, and then TF binding site signals from tumor-specific ATAC-seq mapped regions can be incorporated to our framework.

One limitation of our approach is the multiplicity of inferred effects, which is biologically reasonable but complicates interpretation. Our model also currently makes the assumption that a TF either induces or represses its targets, but some TFs may play either role depending on coordination with co-factors. These limitations may confound the interpretation of inferred TFs with dual activator/repressor roles. Tumor data sets are also a challenging case for regulatory network analysis due to the presence of stromal/immune cells within the tumor and the heterogeneity of cancer cells themselves. However, the PSIONIC framework can be extended modeling of single-cell RNA-seq, as we will report elsewhere.

We used PSIONIC to perform a comprehensive transcriptional network analysis of gynecologic and basal breast cancer tumors. These tumors have not previously been subject to extensive epigenetic or computational analyses, and they all lack accurate predictors of response and treatment strategies for recurrent disease. PSIONIC can identify transcriptional processes that are active across otherwise very different tumors, such as MEF2A activity in the OV, BRCA-BASAL and UCS cohorts. Applying our method to other pan-cancer cohorts such as squamous carcinomas or pediatric cancers might again find biological processes that are activated in a large number of tumor types and provide insight into common regulatory programs in tumors of different origin. We also demonstrated that PSIONIC-predicted activity for TFs in cell line models correlated with sensitivity to TF inhibition, giving a proof-of-principle for the potential therapeutic application of our approach.

Patient-specific inference of TF networks may ultimately enable the development of individualized therapies, aid in understanding mechanisms of drug resistance, and allow the identification of biomarkers of response. We anticipate that computational modeling of transcriptional regulation across different tumor types will emerge as an important tool in precision oncology, aiding in the eventual goal of choosing the best therapeutic option for each individual patient.

## METHODS

### Datasets

RNA-seq data for each of the 5 tumor types were downloaded from TCGA’s Firehose data run (https://confluence.broadinstitute.org/display/GDAC/Dashboard-Stddata). Log 10-transformed RNA-seq RSEM gene expression values were unit-normalized by tumor sample. Cancer cell lines RNA-seq data were downloaded from the CCLE website (http://www.broadinstitute.org/ccle). Log 10-transformed RNA-seq TPM gene expression values were unit-normalized by cell line.

### Sample preparation for ATAC-seq

Sample preparation for ATAC-seq was performed as described previously by Epinomics [26].

### ATAC data analysis

ATAC-seq libraries generated from basal breast (MDA-MB-231, MDA-MB-436) high-grade serous ovarian (OVCAR8, Caov3), uterine carcinosarcoma (JHUCS, SNU685), endometrial endometrioid (AN3-CA, KLE, Ishikawa, RL95-2) and serous carcinoma (ACI-126, ACI-158) cell lines. For each sample, ATAC-seq was performed on two biological replicates.

Starting from fastq files containing ATAC-seq paired-end reads, sequencing adaptors were removed using Trimmomatic [27]. Trimmed reads were mapped to the hg19 human genome using Bowtie2 [28] allowing at most 1 seed mismatch and keeping only uniquely aligned reads. Duplicates were removed using Picard (http://picard.sourceforge.net). For peak calling the read start sites were adjusted (reads aligning to the +/-strand were offset by +4bp/-5bp, respectively) to represent the center of the transposase binding-event, as described in Buenrostro et al. [26].

BigWig files were generated using bamCoverage from the deepTools suite with options --binSize 10 --normalizeTo1 × 2451960000 –ignoreForNormalization chrX. The log_2_-transformed ATAC-seq signal were calculated using bamCompare from deepTools [29]. Resulting normalized BigWig files were used as input to computeMatrix to calculate scores for regions of interest (using either scale-regions or reference-point mode) and visualized using plotHeatmap tool from deepTools.

Peak calling was performed on each cell type individually: first, the reads from different replicates were pooled, and the MACS2.0 peak caller [30] was then used to identify peaks with a permissive threshold (*P* < 2×10^−3^). Finally, IDR was used to identify reproducible peaks for each cell type (IDR < 1×10^−2^). Peaks found reproducibly in each cancer cell subtype were combined to create a genome-wide atlas of accessible chromatin regions. Reproducible peaks from different samples were merged if they overlapped by more than 75%. The atlas of chromatin accessibility across twelve gynecologic and basal breast cancer cell lines contains 282,248 peaks. The number of reproducible peaks for each cell line and number of peaks in each cancer type specific atlas are listed in **Supplementary Table 1**.

We associated each peak to its nearest gene in the human genome using the ChIPpeakAnno package [31]. ATAC-seq peaks located in the body of the transcription unit, together with the 100kb regions upstream of the TSS and downstream of the 3’ end, were assigned to the gene.

Using the MEME [32] curated CisBP [33] TF binding motif reference, we scanned each ATAC-seq tumor type peak atlas with FIMO [34] to find peaks likely to contain each motif (*P*?<?10^−5^). We filtered TFs that were not expressed in at least 50% of samples in at least one of the five tumor types. Further, similarity of predicted target peak sets was measured using the Jaccard index (size of intersection/size of union). If two TFs had a high Jaccard index (greater than 0.5), we looked at the mean Jaccard index of each TF with all other TFs, and we removed the TF with the largest mean Jaccard index. The final set contained 352 motifs.

We created a matrix that defines a candidate set of associations between TFs and target genes: TF binding site identification was used to turn each gene’s set of assigned ATAC peaks into a feature vector of binding signals by assigning the maximum score of each motif across all peaks to a gene.

### Differential peak accessibility

Reads aligning to atlas peak regions were counted using the countOverlaps function of the R packages GenomicAlignments and GenomicRanges [35]. Differential accessibility of these peaks was then calculated for all pairwise comparisons of cancer types using DESeq2 [36].

### Motifs underlying differential accessibility in cell lines

The shift in the cumulative distribution of chromatin accessibility changes (log2 fold change) of the subset of the atlas occupied by each TF, compared to that of the background atlas, was measured by a one-sided Kolmogorov-Smirnov (K-S) test in either direction. The foreground occurrence is the number of peaks containing a particular TF motif within the group of 5000 differentially open or 5000 differentially closed peaks according to log2 fold change read counts. The background occurrence is the number of peaks containing a particular TF motif among all the differentially accessible peaks.

### Multitask learning

For multitask learning, we trained regression models jointly for all tumors using *grouping and overlap in multitask learning* (GO-MTL) [7]. In this approach, prediction of each gene expression **y**_*t*_ is considered one task, and we wish to solve T tasks jointly so that information is “shared” between them. Let **X** be the data matrix of size d × N where each row represents a gene and each column is a motif hit score representing the target genes of a TF.

Briefly, we assume there are k (< T) latent basis tasks and each observed task can be represented as linear combination of a subset of these basis tasks. This assumption enables us to write the weight matrix as **W = LS**, where **L** is a matrix of size d x k with each column representing a latent task, and **S** is a matrix of size k x T containing the weights of linear combination for each task. **Ls**_t_ gives the predictor **w**_t_ for task t, where **s**_t_ is t-th column of matrix **S**. The matrix **L** captures the predictive structure of the tasks and the grouping structure is determined by matrix **S**. Tasks that have same sparsity pattern can be seen as belonging to the same group, while tasks whose sparsity patterns are orthogonal to each other can be seen as belonging to different groups. The partial sharing of latent basis tasks allows us to do away with the concept of disjoint groups. Any task that does not share latent bases with any other task in the pool can be seen as outlier task. Our learning cost function takes the following form:

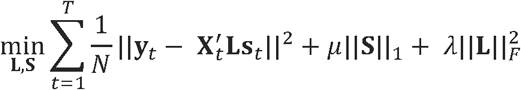

The parameter µ controls the sparsity in **S**. The penalty on the Frobenius norm of **L** regularizes the predictor weights to have low ℓ_2_ norm and avoids overfitting.

To assess single task learning (STL) performance, we trained ridge regression model s for each tumor (task) independently. We fit the ridge regression models using the SLEP MATLAB package and evaluated performance on held-out genes.

### *In vitro* drug sensitivity analysis

Detailed information on cell culture media is provided in the **Supplementary Table 4**. All cell lines were cultured under standard conditions at 37°C and 5% CO_2_. Cells were plated at 10-20% confluency (with the exception of JHUCS-1, MDA-MB-436, and RL-95 which were plated at ∼50%) in 24-well plates in complete medium, and incubated inside an IncuCyte ZOOM system (Essen BioScience, Inc., MI, USA). The following day (22-24 hours later), cells were exposed to LOR-253 (MedChemExpress, NJ, USA) at 0, 50, 250, or 1250 nM. To monitor cell growth, phase contrast images of the cell cultures in the presence or absence of the drug were captured automatically at 2-hour intervals for up to 36 hours, and occupied area of the cells (% confluency) was calculated using the IncuCyte image analysis software. We analyzed drug response data using a recently developed “growth rate inhibition” (GR) metric that corrects for differences in cell proliferation rates [37].

### Immunohistochemical staining

FFPE 4 µm sections from patient tissues were deparaffinized and fixed in methanol prior to antigen retrieval in heated Citrate Buffer (pH 6.0, Poly Scientific R&D Corp.) at 95° C for 15 cubes used were DAPI (cubes used were DAPI (cubes used were DAPI (cubes used were DAPI (min (EZ Retriever microwave, BioGenex). Sections underwent two rounds of staining, each including a peroxidase block, protein block, primary and secondary antibody incubation, and fluorophore binding. Between both staining rounds, samples underwent antigen retrieval. Endogenous peroxidase was blocked with Hydrogen peroxide solution (Millipore Sigma) 3% for 10 minutes. Protein blocking was performed using PBS-Tween 3% BSA 1% donkey serum (Millipore Sigma) for 30 minutes. Each section was stained first with ETV6 (1:100, polyclonal, Millipore Sigma), followed by cytokeratin (OSCAR) (1:100, Cell Marque, Millipore Sigma). A broad-spectrum HRP secondary antibody, Superpicture Polymer Detection Kit (Thermo Fisher) was used. Fluorophores Opal 540 (ETV6) and Opal 620 (Cytokeratin) were covalently bound using tyramide signal amplification (Perkin Elmer). Nuclei were stained with 4,6-diamidino-2-phenylindole (DAPI) (Thermo Fisher) and mounted with ProLong Diamond Antifade mountant (Thermo Fisher).

### Imaging and analysis

Slides were imaged using the Vectra Multispectral Imaging system version 2 (Perkin Elmer). The filter cubes used were DAPI (440–680 nm), FITC (520 nm-680 nm) and Texas Red (580–700 nm). A spectral library with the emitting spectral peaks of the fluorophores used was created with inForm 2.1 image analysis software (Perkin Elmer) by using multispectral images obtained from single-stained slides for each marker and associated fluorophore. The spectral library allows for spectral unmixing of each multispectral image. Unmixed images were then subjected to an active learning algorithm using inForm, determining the tumor and stromal components, and the nuclear and cytoplasmic compartment in each image. Subsequently, 2-bin scoring was used to determine positive or negative expression of ETV6 in every tissue and cellular compartment.

## Software and data availability

The ATAC-seq data have been deposited in the Gene Expression Omnibus accession number GSE114964. The software for PSIONIC is available from https://sites.google.com/view/haticeulkuosmanbeyoglu/software.

## Acknowledgements

We would like to thank John I. Risinger for sending us the ACI158 and ACI126 cell lines. The results published here are in whole or part based on data generated by The Cancer Genome Atlas project established by the NCI and NHGRI (accession number: phs000178.v7p6). Information about TCGA and the investigators and institutions that constitute the TCGA research network can be found at http://cancergenome.nih.gov/. This work was supported by NCI R21 award CA205819. H.U.O. is supported by NCI K99 award CA207871.

## Author Contributions

H.U.O and C.S.L. conceived and designed the study, and developed the computational framework. H.U.O carried out the model training and computational validation. H.U.O and C.S.L. analyzed data and wrote the manuscript. F.S. performed the experimental validation for *In vitro* drug sensitivity and under supervision of. G.C. and helped to write the experimental validation section. A.R.V. performed the Immunohistochemical staining under supervision of S.C.M. and helped to write the experimental validation section. P.J. and D.A.L. assisted with study design.

## SUPPLEMENTARY FIGURE LEGENDS

**Supplementary Figure 1:**
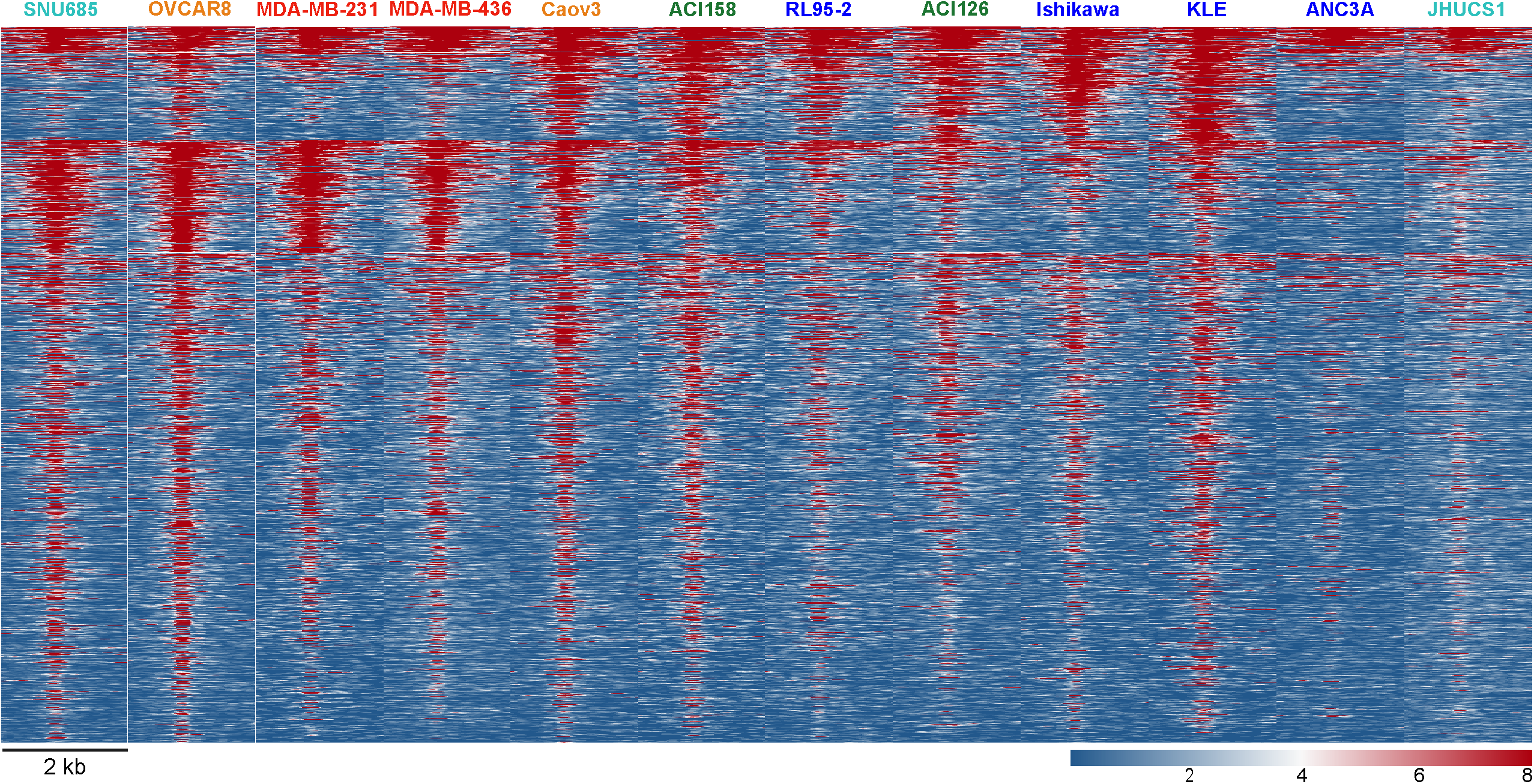
Chromatin accessibility heat map grouped by differential accessibility patterns. Each row represents one of 40,000 selected peaks (differentially accessible between at least one histologic type comparison; FDR < 0.0001, log2(fold-change; FC) > 3).

**Supplementary Figure 2:**
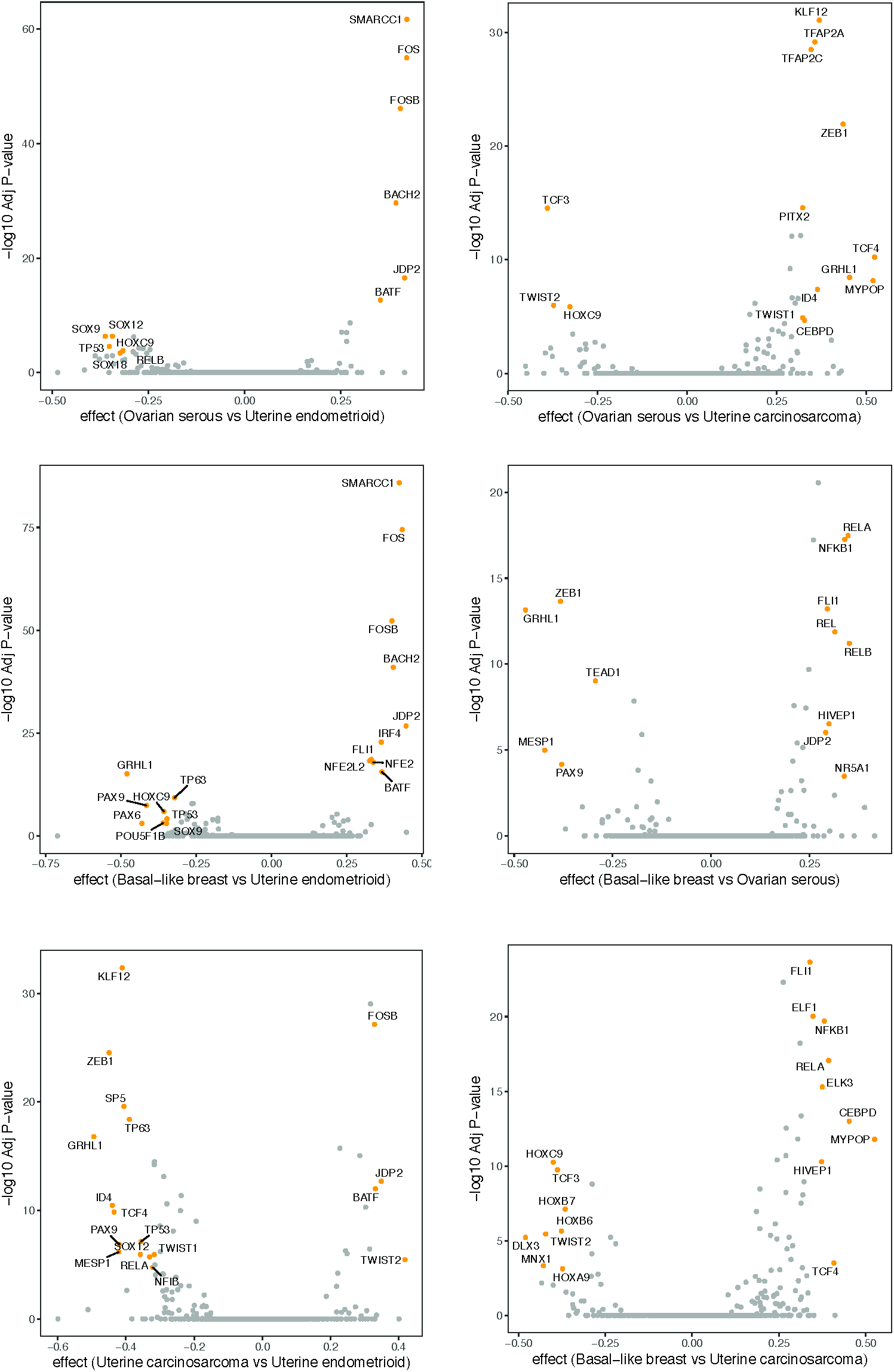
Motifs underlying differential accessibility in cell lines. Volcano plot showing effect size versus –log10(Adjusted P-value) for that comparison.

**Supplementary Figure 3:**
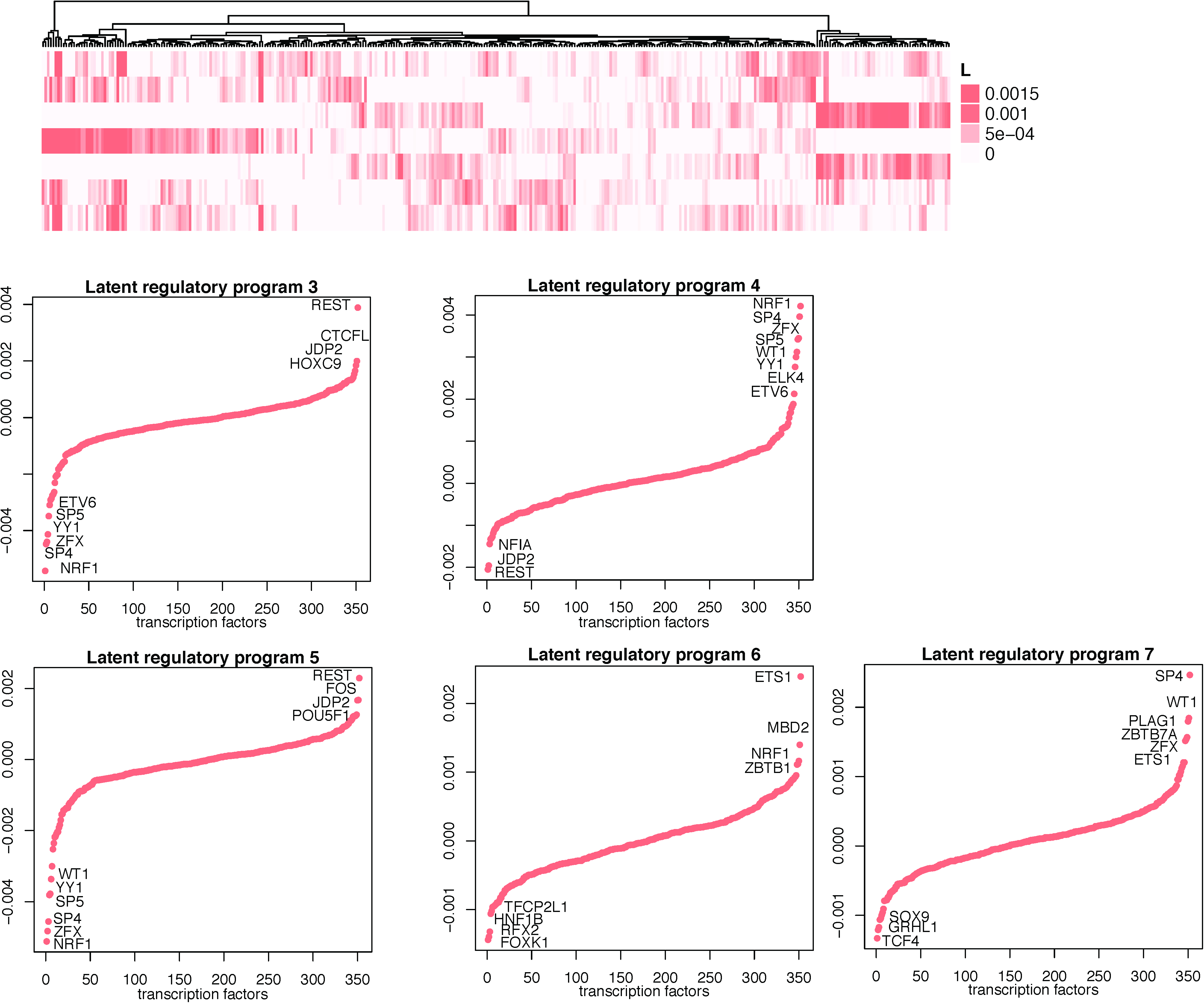
Heatmap of latent regulatory programs/tasks. Regulators are ranked based on increase in magnitude of coefficients.

**Supplementary Figure 4:**
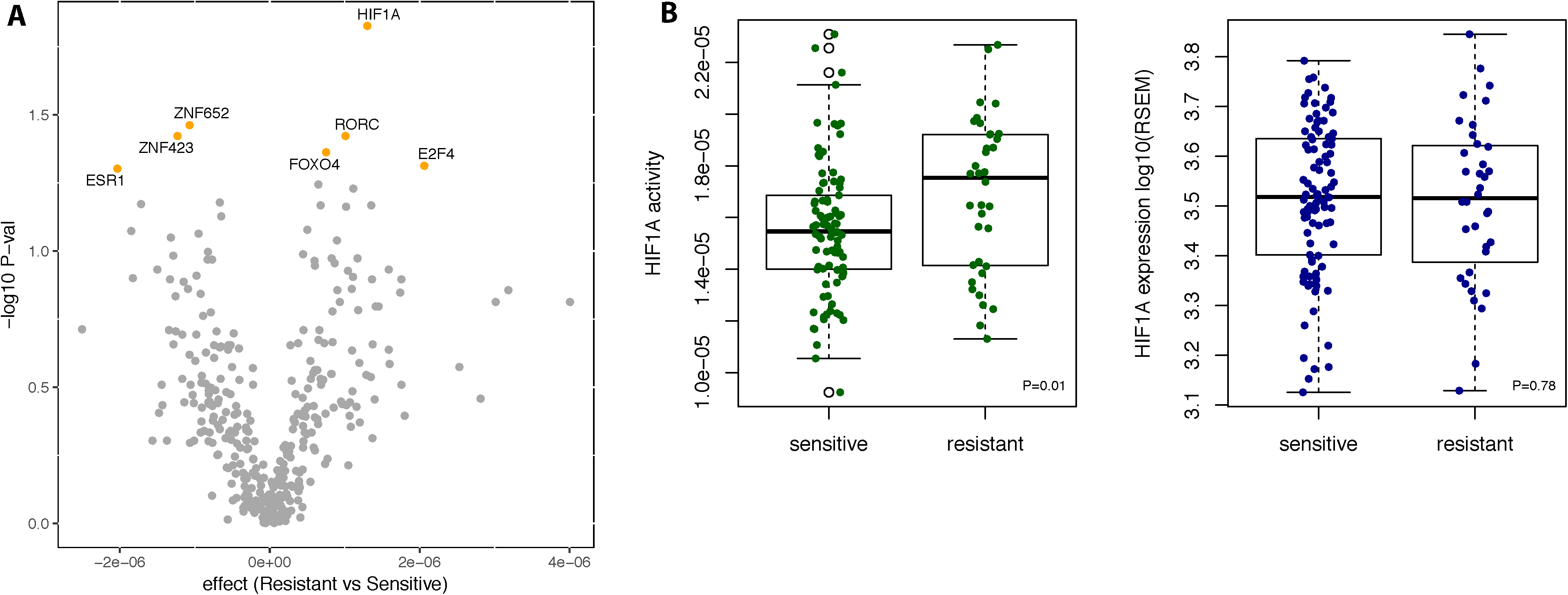
t-SNE projections of mean centered inferred TF activities for TCGA patients (as donated with squares) and CCLE cell lines (as donated with plus sign).

**Supplementary Figure 5:**
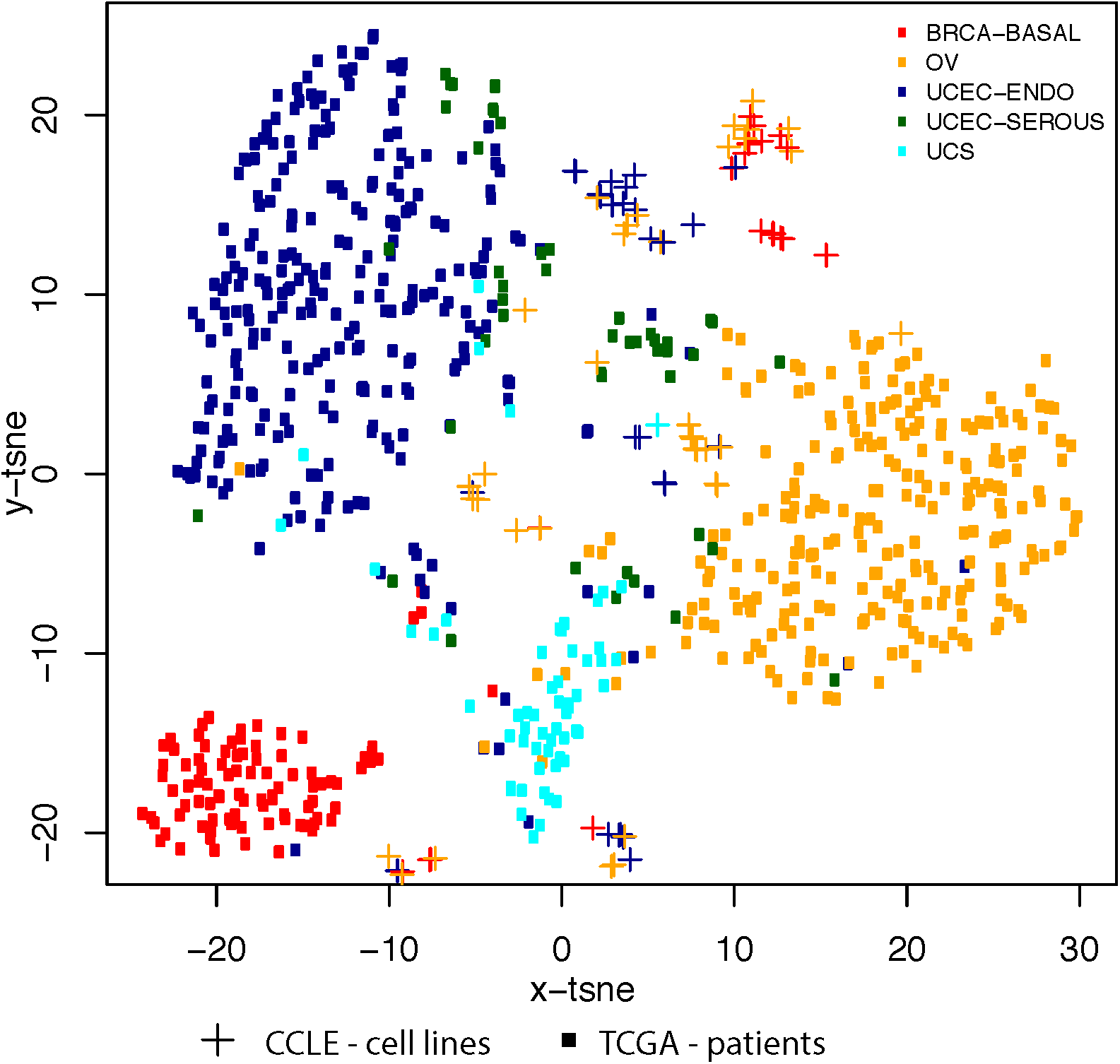
**(A)** The mean inferred TF activity difference in platinum resistant and sensitive patients is plotted on the *x* axis, and p values from *t*-test is plotted on the *y* axis (–log10 scale) for OV cohort. TFs associated with response (p-value <0.02) are coloured in orange. **(B)** HIF1A differential inferred TF activities in platinum resistant and sensitive patients tumors (*P*<10^−1^, Wilcoxon’ s rank-sum test). Importantly, HIF1A gene expression is not statistically significantly different in gene expression level (right side).

## SUPPLEMENTARY TABLES

**Supplementary Table 1:** Summary of ATAC-seq data.

| Cell line | Histological subtype | Reproducible peaks | Atlas peaks |
| --- | --- | --- | --- |
| MDA-MB-231 | Basal breast | 69,136 | 102,981 |
| MDA-MB-436 |  | 77,999 |  |
| CAOV3 | Ovarian adenocarcinoma | 102,048 | 153,681 |
| OVCAR8 |  | 110,600 |  |
| JHUCS | Uterine carcinosarcoma | 50,483 | 135,948 |
| SNU685 |  | 121,132 |  |
| AN3CA | Endometrial endometrioid adenocarcinoma | 54,668 | 147,024 |
| Ishikawa |  | 68,207 |  |
| KLE |  | 83,824 |  |
| RL95_2 |  | 50,221 |  |
| ACI-126 | Endometrial serous adenocarcinoma | 56,175 | 93,976 |
| ACI-158 |  | 69,918 |  |

**Supplementary Table 4:**
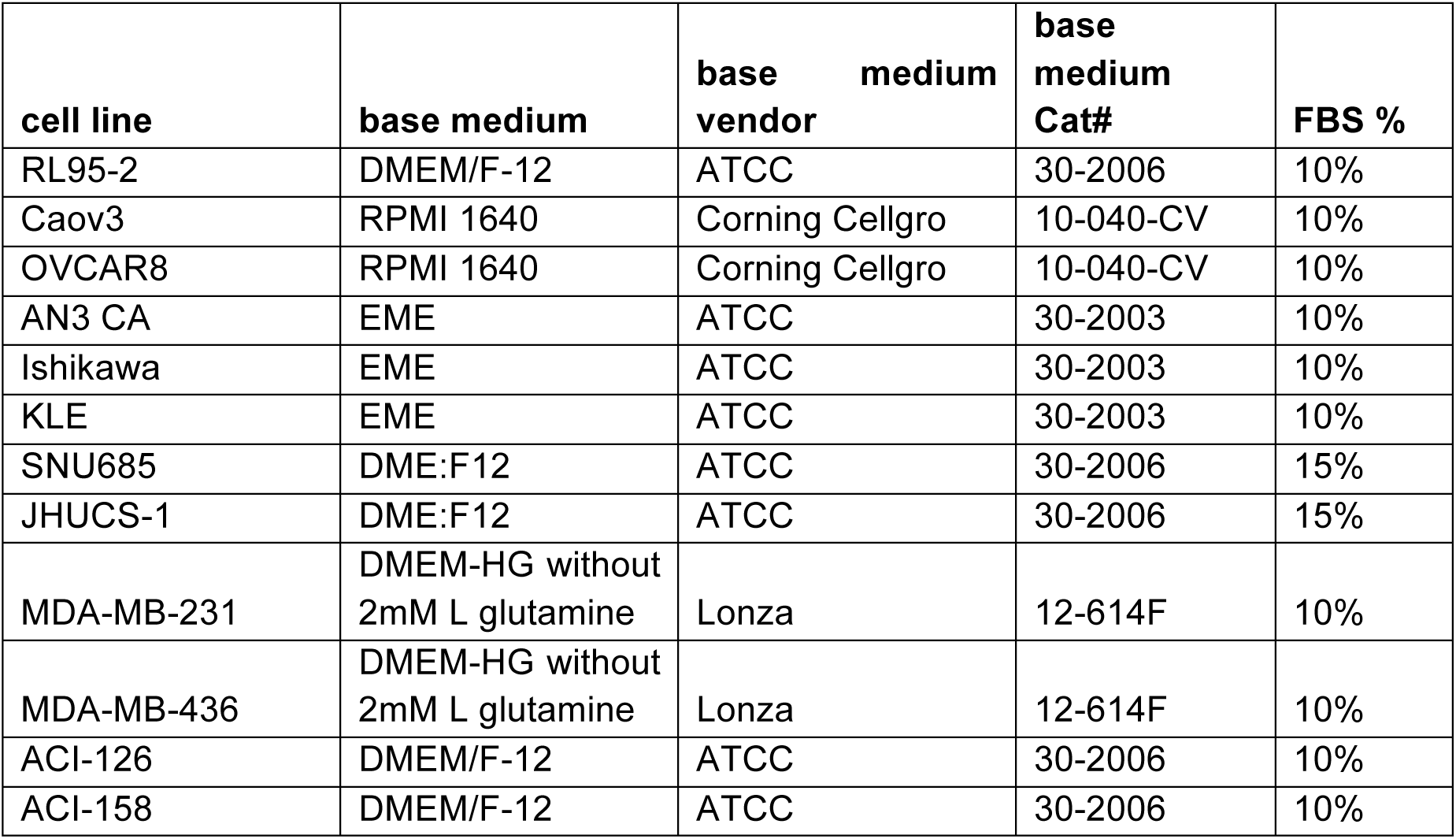
Cell culture media

